# Local Metabolism of Ethanol to Acetic Acid/Acetate in the Central Nucleus of Amygdala Elicits Sympathoexcitatory Responses through Activation of NMDAR in Sprague Dawley Rats

**DOI:** 10.1101/2020.07.20.212597

**Authors:** Andrew D. Chapp, Michael J. Huber, Andréa R. Collins, Kyle M. Driscoll, Jessica E. Behnke, Robert A. Larson, Zhiying Shan, Li. Zhang, Qing-Hui Chen

**Author notes:** Correspondence to: Qing-Hui Chen, Ph.D. Department of Kinesiology and Integrative Physiology, Michigan Technological University. SDC, 1400 Townsend Drive, Houghton, MI 49931, Andrew D. Chapp, Ph.D. Department of Neuroscience, University of Minnesota-Twin Cities, 321 Church St SE, Minneapolis, MN 55455.

## Abstract

Binge alcohol consumption elicits robust sympathoexcitation and excitatory neuronal output. However, the central mechanism that mediates these effects remains elusive. We investigated the effects of ethanol metabolism within the central nucleus of the amygdala (CeA) on sympathoexcitation, and elucidated the role of acetate in these excitatory responses. *In vivo* arterial blood pressure, heart rate and sympathetic nerve activity responses to CeA microinjected ethanol or acetate with appropriate inhibitors/antagonists were tracked. *In vitro* whole-cell electrophysiology recording responses to acetate in CeA neurons with axon projecting to the rostral ventrolateral medulla (CeA-RVLM) were investigated, and cytosolic calcium responses in primary neuronal cultures were quantified. We demonstrate that in Sprague Dawley rats, local brain metabolism of ethanol in the CeA to acetic acid/acetate elicits sympathoexcitatory responses *in vivo* through activation of NMDA receptor (NMDAR). Alcohol dehydrogenase or aldehyde dehydrogenase inhibition using fomepizole or cyanamide and NMDAR antagonism using AP5 or memantine blunted these effects. Whole-cell patch-clamp recordings in brain slices containing autonomic CeA-RVLM neurons revealed a dose-dependent increase in neuronal excitability in response to acetate. NMDAR antagonists suppressed the acetate-induced increase in CeA-RVLM neuronal excitability, and memantine suppressed the direct activation of NMDAR-mediated inward currents by acetate in brain slices. We observed that acetate increased cytosolic Ca^2+^ in a time-dependent manner in primary neuronal cell cultures. The acetate enhancement of calcium signaling was abolished by memantine. These findings suggest that within the CeA, ethanol is sympathoexcitatory through local brain metabolism, which generates acetic acid/acetate leading to activation of NMDAR.

**NEW AND NOTEWORTHY:** Brain ethanol metabolism to acetic acid (vinegar)/acetate causes activation of N-methyl-D-aspartate receptors (NMDARs) in the central nucleus of the amygdala and elicits sympathoexcitatory responses. This excitatory mechanism is opposite to the inhibitory effects of ethanol at NMDAR. Understanding the active compounds that arise from ethanol metabolism, and the molecular mechanisms by which they influence alcohol reward and cardiovascular function, may be beneficial in developing targeted intervention strategies for both alcohol use disorder and its cardiovascular sequelae.

**Graphical Figure:** Proposed mechanisms for ethanol and acetate induced increases in sympathoexcitation within the central nucleus of the amygdala (CeA). Abbreviations: Acetic acid (HOAc), acetate (^-^OAc), ADH (alcohol dehydrogenase), ALDH (aldehyde dehydrogenase), BBB (blood brain barrier), FOM (fomepizole), CYAN (cyanamide), CYP450 (cytochrome P450), IML (intermediolateral nucleus), RVLM (rostral ventrolateral medulla), SNA (sympathetic nerve activity).

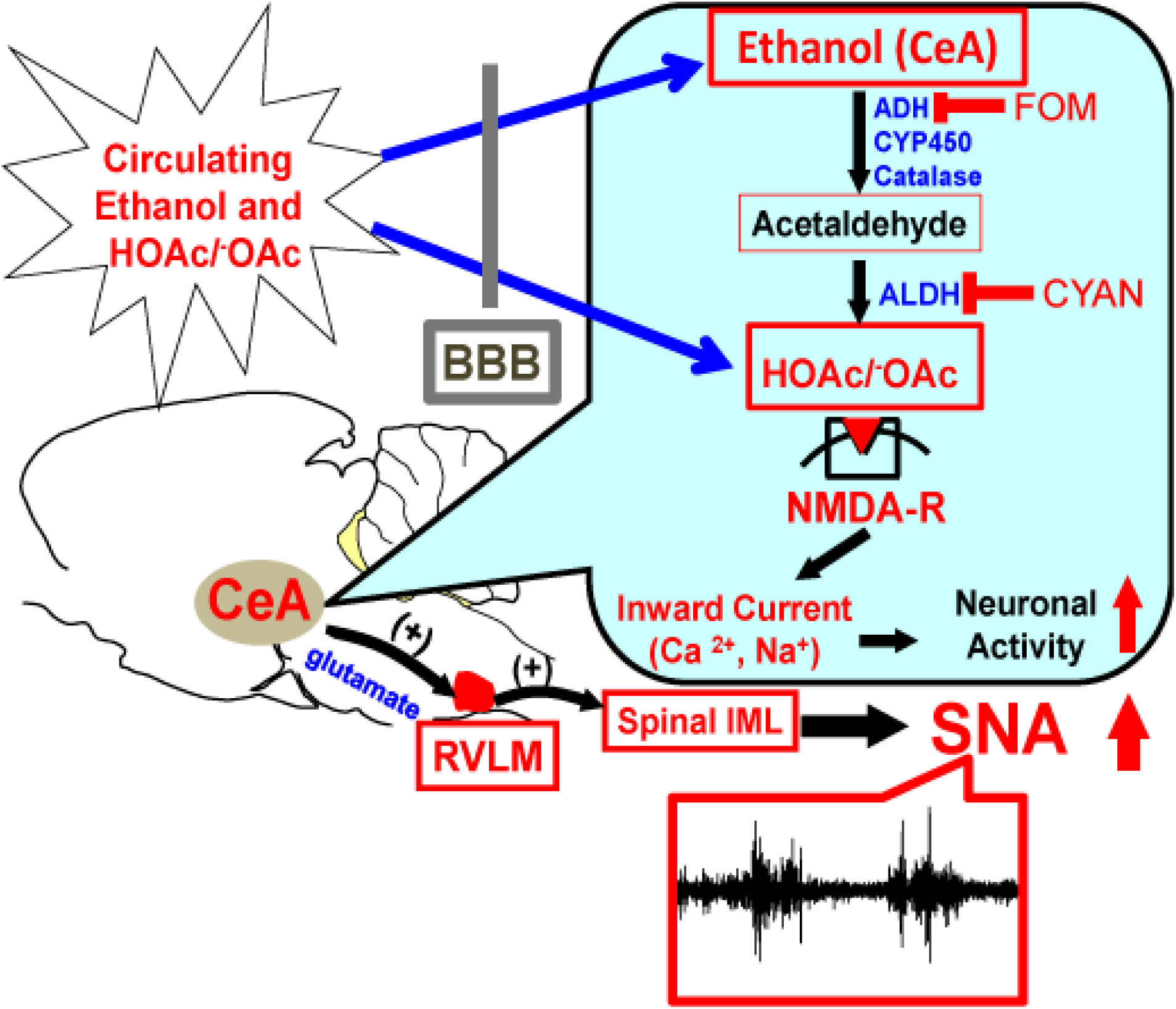

## INTRODUCTION

Alcohol is one of the most widely used and abused recreational substances globally. Ethanol metabolism generates acetic acid, which is the main component of vinegar and is produced in large quantities following ethanol ingestion. This leads to significantly elevated serum and cerebrospinal fluid (CSF) acetate concentrations of around 1-3 mM (1, 2) and 2-5 mM (1, 2), respectively. Concentrations of acetic acid/acetate in the circulation and CSF are reported to remain elevated long after ethanol has cleared the body (2). It is therefore reasonable to suspect that acetic acid/acetate may be a player in the altered neuronal function following ethanol consumption and metabolism.

Precisely how ethanol produces excitatory effects in the central nervous system (CNS) is largely unknown. On the one hand, increases in dopamine release from the ventral tegmental area (3) facilitate reward and possible excitatory actions on neurons post ethanol ingestion. However, this is partially contradicted by ethanol itself inhibiting NMDAR in slice recordings (4–6). Secondly, studies involving NMDAR inhibition by ethanol are commonly done in brain slice preparation (4, 5, 7) with large circulating volumes of ethanol relative to small tissue samples, and the effects of in-slice metabolism of ethanol has yet to be reported. What is known is that ethanol consumption contributes to excitatory actions originating within the CNS (3, 8) with some positive modulatory action on NMDAR (9). Excitation is augmented following ethanol clearance, which can lead to hangover symptoms (10, 11). Acetate has been implicated in this response (12). Unfortunately, literature investigating the influence of acetic acid/acetate on neuronal function remains scarce.

Our lab has been pursuing a line of thought that acetic acid/acetate is responsible for many of the excitatory effects of ethanol metabolism on neurons. We have previously reported that microinjection of ethanol and acetate into the central nucleus of amygdala (CeA), an area containing autonomic neurons with axons projecting to the rostral ventrolateral medulla (CeA-RVLM), induced sympathoexcitation and pressor responses mediated by local activation of NMDAR (13). Similarly, we have demonstrated that pathophysiological concentrations of acetate from ethanol metabolism are cytotoxic in dopaminergic-like PC12 cells, at least partially through NMDAR activation (14) and acetic acid increases excitability in medium spiny neurons of the nucleus accumbens shell (15). Given these parallel and converging sets of results, we hypothesized that inhibition of ethanol metabolism (Figure. 1A) may attenuate the ethanol sympathoexcitatory response by preventing the generation of acetic acid/acetate which we have found to activate NMDAR. Similarly, we hypothesized that in brain slice recordings containing CeA-RVLM neurons, bath application of acetate would increase neuronal excitability through activation of NMDAR.

**Figure 1.**
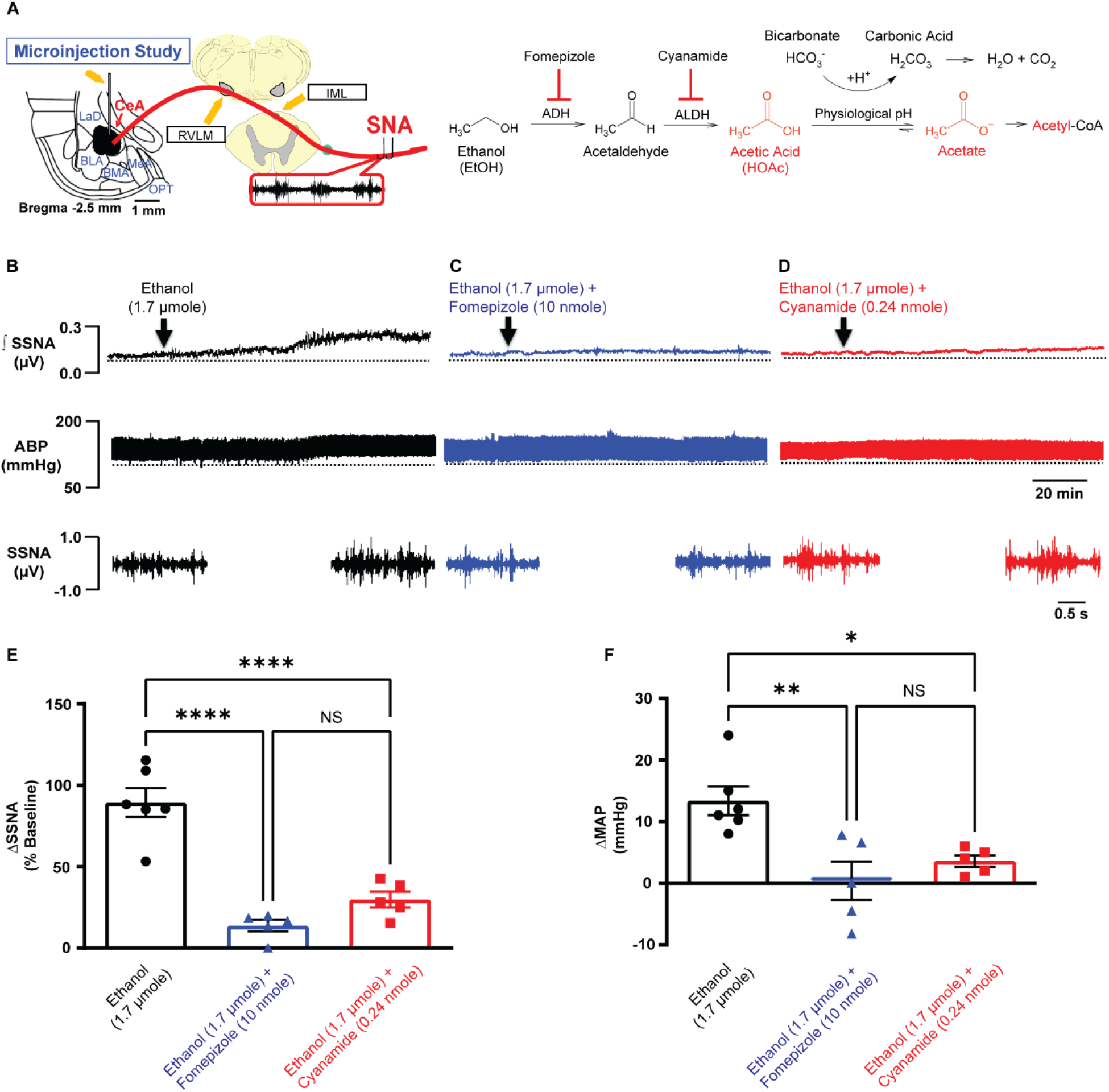
Local CeA brain metabolism of ethanol is sympathoexcitatory. **(A)** Graphical depection of CeA microinjection study for sympathetic nerve recordings (left). Metabolic pathway of alcohol metabolism (right). **(B)** Representative raw traces displaying SSNA and arterial blood pressure (ABP) from CeA microinjected ethanol. **(C)** Representative raw traces displaying SSNA and arterial blood pressure (ABP) from CeA microinjected ethanol + fomepizole. **(D)** Representative raw traces displaying SSNA and arterial blood pressure (ABP) from CeA microinjected ethanol + cyanamide. **(E)** Summary data of SSNA for ethanol (black), ethanol + fomepizole (blue) and ethanol + cyanamide (red) (****p<0.0001). **(F)** Summary data of change in mean aterial pressure (MAP) for ethanol (black), ethanol + fomepizole (blue) and ethanol & cyanamide (red), (n=number of rats) (**p=0.0043, *p=0.029). Abbreviations: ADH, alcohol dehydrogenase; ALDH, aldehyde dehydrogenase).

## MATERIALS AND METHODS

### Animals

Male Sprague-Dawley rats purchased from Charles River Labs (Wilmington, MA, USA) were individually housed in a temperature-controlled room (22-23 ° C) with a 12:12 h light-dark cycle. Chow and tap water were available *ad libitum* unless otherwise noted. All experimental and surgical procedures were carried out under the guidelines of the National Institutes of Health Guide for the Care and Use of Laboratory Animals with the approval of the Institutional Animal Care and Use Committee at Michigan Technological University.

### *In vivo* whole animal recordings of arterial blood pressure (ABP), splanchnic sympathetic nerve activity (SSNA) and lumbar sympathetic nerve activity (LSNA)

Animal surgery was performed following the previously described protocol (16). On the day of the experiment, rats were anesthetized with an intraperitoneal injection containing a mixture of α-chloralose (80 mg/kg) and urethane (800 mg/kg). Adequate depth of anesthesia was assessed before surgery by the absence of the pedal withdrawal reflex and corneal reflexes. Animals were instrumented with an arterial catheter inserted into the aorta through a femoral artery. The catheter was connected to a pressure transducer to measure arterial blood pressure. Heart rate (HR) was obtained from the R-wave of the electrocardiogram (ECG) (lead I). A catheter was also placed in the left femoral vein to administer drugs. After tracheal cannulation, rats were paralyzed with gallamine triethiodide (25 mg·kg-1 ·h-1 iv) and artificially ventilated with oxygen-enriched room air. After paralysis, adequate depth of anesthesia was determined by lack of pressor responses to noxious foot pinch. Supplemental doses of anesthesia equal to 10% of the initial dose were given when needed. End-tidal PCO_2_ was continuously monitored and maintained within normal limits (35–40 mmHg) by adjusting ventilation rate (70–80 breaths/min) and/or tidal volume (2.0–3.0 ml). Body temperature was held at 37°C with a water-circulating pad.

SSNA and LSNA recordings were performed according to previously described protocols (13, 16, 17). With the use of a left flank incision, a left lumbar and postganglionic splanchnic sympathetic nerve bundle was isolated from surrounding tissue and mounted on a stainless steel wire electrode (0.127-mm OD; A-M Systems), and covered with a silicon-based impression material (Coltene, Light Body) to insulate the recording from body fluids. The recorded signal was directed to an AC amplifier (P511; Grass Technologies) equipped with half-amplitude filters (band pass: 100-1,000 Hz) and a 60-Hz notch filter. The processed signal was rectified, integrated (10-ms time constant), and digitized at a frequency of 5,000 Hz using a 1401 Micro3 analog-to-digital converter and Spike 2 software (7.04 version; Cambridge Electronic Design, Cambridge, UK). The background noise was determined by a bolus injection of hexamethonium (30 mg/kg iv), a ganglionic blocker, at the end of the experiment and was subtracted from all the integrated values of sympathetic nerve activity.

### CeA microinjection

CeA injections were performed as previously described (13). Animals were placed in a stereotaxic head frame, and the skull was leveled between bregma and lambda. A section of skull was removed so that a single-barreled glass microinjector pipette could be lowered vertically into the CeA. The following stereotaxic coordinates were used: CeA, caudal to bregma, −2.4 ~ −2.5 mm; lateral to midline, 4.8~5.0 mm; and ventral to dura, 7.7~8.0 mm. All animals were allowed to stabilize at least 2 h following surgery. To test whether inhibition of ADH and ALDH prevented ethanol elicited sympathoexcitatory responses, ethanol (1.7 μmol) or a cocktail of ethanol (1.7 μmol) and fomepizole (10 nmol) or ethanol (1.7 μmol) and cyanamide (0.24 nmol) were microinjected into the CeA in separate groups of animals. Similarly, the effects of acetate and the involvement of NMDAR on acetate evoked sympathoexcitatory response were also explored through CeA injection of acetate (0.1 μmol and 0.20 μmol) and a cocktail containing acetate (0.20 μmol) and memantine (3.0 nmol), or acetate (0.2 μmol) and AP5 (3.0 nmol) (13, 18) in separate groups of animals. Likewise, to verify lack of involvement of cyanamide in the acetate response, a cocktail of acetate (0.20 μmol) and cyanamide (0.24 nmol) was microinjected in separate groups of animals. All compounds were microinjected in a volume of 100 nL with a pneumatic pump (WPI). The volume of each injection was determined by measuring the movement of the fluid meniscus within the microinjector pipette using a dissecting microscope equipped with an eyepiece reticule. At the end of each experiment, Chicago blue dye solution (2% in saline, 100 nL) was injected into the CeA to mark the site of each injection. Brains were removed and post fixed for 5 days at room temperature in 4% paraformaldehyde. Brain coronal sections containing the CeA were cut, and microinjection sites were identified under light microscopy. Rats with injection site(s) not inside the CeA were excluded from data analysis.

### Retrograde labeling

Retrograde labeling of CeA-RVLM neurons(13): Male Sprague-Dawely rats (350-450 g) were retrograde labeled from the RVLM to the CeA. Rats were anesthetized with an intraperitoneal injection (i.p.) of sodium pentobarbital (50 mg/kg), placed in a stereotaxic frame, leveled between bregma and lambda and the skull exposed with a midline incision. A small, unilateral burr hole was drilled to expose the cerebellum in the caudal section of the skull and a glass micropipette lowered into the pressor region of the RVLM (coordinates: −12.7 mm caudal to bregma, 1.8 mm lateral to midline and 8.9 mm below the skull) (19). 100 nL fluospheres (Life Technologies) was injected into the RVLM using a pneumatic pump (WPI) and left in place for approximately 5 minutes to prevent tracer backflow. The midline incision was closed, and the rats were administered a subcutaneous (s.c.) injection of penicillin G (30,000 units) and Metacam (1 mg/kg, NSAID) in 0.5 mL saline for three days post-surgery to prevent infection and reduce pain.

### Whole-cell patch clamp recordings

Five to seven days post labeling, rats were anesthetized with isoflurane (3% in O_2_) and decapitated. The brain was rapidly removed and chilled in ice cold cutting solution, containing (in mM): 206 sucrose, 2 KCl, 2 MgSO_4_, 1.25 NaH_2_PO_4_, 26 NaHCO_3_, 1 CaCl_2_, 1 MgCl_2_, 10 d-glucose, and 0.4 ascorbic acid, osmolarity 295–302 mosmol L^-1^ measured with an osmometer (Wescor), pH 7.3-7.4, continuously gassed with 95:5 O_2_:CO_2_ to maintain pH and pO_2_. A brain block was cut including the CeA region and affixed to a vibrating microtome (Leica VT 1000S; Leica, Nussloch, Germany). Coronal sections of 250 μm thickness were cut, and the slices transferred to a holding container of artificial cerebral spinal fluid (ACSF) maintained at 30 °C, continuously gassed with 95:5 O_2_:CO_2_, containing (in mM): 125 NaCl, 2 KCl, 2 MgSO_4_, 1.25 NaH_2_PO_4_, 26 NaHCO_3_, 2 CaCl_2_, 10 d-glucose, and 0.4 ascorbic acid (osmolality: 295-302 mosmol L^-1^; pH 7.3-7.4) and allowed to recover for 1 hr. Following recovery, slices were transferred to a glass-bottomed recording chamber and viewed through an upright microscope (E600FN, Nikon) equipped with DIC optics, epi-fluorescence, an infrared (IR) filter and an IR-sensitive video camera (C2400, Hamamatsu, Bridgewater, NJ).

Slices transferred to the glass-bottomed recording chamber were continuously perfused with ACSF, gassed with 95:5 O_2_:CO_2_, maintained at 30 °C and circulated at a flow of 2 mL min^-1^. Patch electrodes were pulled (Flaming/Brown P-97, Sutter Instrument, Novato, CA) from borosilicate glass capillaries and polished to a tip resistance of 4–8 MΩ. Electrodes were filled with a solution containing (in mM) 135 K-gluconate, 10 HEPES, 0.1 EGTA, 1.0 MgCl_2_, 1.0 NaCl, 2.0 Na_2_ATP, and 0.5 Na_2_GTP (osmolality: 280–285 mosmol L^-1^; pH 7.3). Once a GΩ seal was obtained, slight suction was applied to break into whole-cell configuration. The cell was allowed to stabilize; this was determined by monitoring capacitance, membrane resistance, access resistance and resting membrane potential (V_m_) (19). Records were not corrected for a liquid junction potential of −15 mV. Cells that met the following criteria were included in the analysis: action potential amplitude ≥50 mV from threshold to peak, input resistance (*R*_input_) >0.3 GΩ (determined by injection of −20 pA from a holding potential of −80 mV), resting *V*_m_ negative to −50 mV, and <20% change in series resistance during the recording. Recordings were made using an Axopatch 200B amplifier and pCLAMP software (v10.2, Axon Instruments, Union City, CA). Signals were filtered at 1 kHz, digitized at 10 kHz (Digidata 1400A, Axon Instruments), and saved on a computer for off-line analysis. To study the mAHP as we have previously reported,(19) recordings were made from a potential of −60 mV, and +150 pA current injection (500 ms) were made followed by a return of *V*_m_ to −60 mV for 5 s.

The effect of acetate on the excitability of CeA-RVLM neurons and the role of NMDAR (current-clamp): With V_m_ adjusted to −80 mV by continuous negative current injection, a series of square-wave current injections was delivered in steps of +25 pA, each for a duration of 800 ms. To determine the action potential voltage threshold (V_t_ or rheobase) and depolarizing *R*_input_ below *V*_t_, ramp current injections (0.2 pA/ms, 1,000 ms) were made from a potential of −80 mV. Square-wave and ramp current injections were made in the same neurons. Effects of acetate on neuronal excitability were determined by comparing current evoked spike frequency responses under control conditions with responses recorded from neurons exposed to bath application of acetate. To determine the dose-dependent response, the effect of various concentration of acetate (0.0, 3.75, 7.5, 37.5 and 75.0 mM) on the excitability was determined in separate groups of neurons, respectively. To test whether the activation of NMDAR was involved in the increased neuronal excitability elicited by acetate, the effect of bath application of AP5 (60 μM, NMDAR competitive blocker) or memantine (30 μM, NMDAR pore blocker), on the excitability was determined in the presence of acetate (37.5 mM) in a separate group of neurons. To determine the slope of the current injection response (another measure of how excitable neurons are over a range of current injection), a linear regression was fit to each treatment group (control, NaOAc, NaOAc & AP5 and NaOAc & memantine), between 0 and 250 pA. The slope of each current injection response (gain) ± SEM were compared across groups.

Acetate evoked inward current and the role of NMDAR (voltage-clamp) in CeA-RVLM neurons: Brain slices were continuously perfused with modified, Mg^2+^ free ACSF containing (in mM): 127 NaCl, 2 KCl, 1.25 NaH_2_PO_4_, 26 NaHCO_3_, 2 CaCl_2_, 10 d-glucose, and 0.4 ascorbic acid (osmolality: 295–302 mosmol L^-1^; pH 7.3-7.4), gassed with 95:5 O_2_:CO_2_, maintained at 30 °C and circulated at a flow of 2 mL min^-1^. Tetrodotoxin (TTX, voltage gated sodium channel blocker, 0.5 μM) and picrotoxin (GABA-A blocker, 100 μM) were added into the circulating extracellular, Mg^2+^ free ACSF. Cells were voltage clamped at V_m_ = −80 mV and allowed to stabilize by monitoring baseline current. Once cells were stable and baseline recording in the absence of acetate ~1.5 min was observed, acetate (37.5 mM) was added to the circulating bath and recorded until a plateau was observed > 15 min. At this point, washout and recovery was observed. To test whether the activation of NMDAR was involved in the inward current evoked by acetate, the effect of memantine (30 μM) or AP5 (60 μM) on the inward current was determined. In separate groups of neurons, acetate (37.5 mM) and memantine (30 μM) or acetate (37.5 mM) and AP5 (60μM) were added to the circulating bath and the inward current was observed. To exclude the possibility that the inward current evoked by sodium acetate was due to the increase in extracellular sodium concentration, equimolar sodium gluconate (37.5 mM) was substituted for sodium acetate and the inward current was recorded.

Note that current- and voltage-clamp recordings were made from separated groups of CeA-RVLM neurons. Passive membrane properties (capacitance, resting membrane potential and hyperpolarizing input resistance) to each drug treatment protocol are reported in table 1.

**Table 1.** Membrane properties of CeA-RVLM neurons

| Group | Neurons<br>(n) | V<sub>m</sub><br>(mV) | C<sub>m</sub><br>(pF) | HR<sub>input</sub><br>(GΩ) | V<sub>t</sub><br>(mV) | Rats<br>(n) |
| --- | --- | --- | --- | --- | --- | --- |
| Control | 7 | -58.0 ± 3.0 | 47.4 ± 2.5 | 0.47 ± 0.07 | -43.8 ± 1.8 | 5 |
| NaOAc<br>(7.5 mM) | 6 | -59.5 ± 2.6 | 45.3 ± 4.5 | 0.53 ± 0.08 | -42.4 ± 2.7 | 3 |
| NaOAc<br>(37.5 mM) | 8 | -62.3 ± 2.2 | 34.6 ± 2.5 | 0.58 ± 0.03 | -49.3 ± 1.8 | 3 |
| NaOAc<br>(37.5 mM)<br>+<br>AP5<br>(60 μM) | 7 | -67.9 ± 4.3 | 36.3 ± 2.2 | 0.45 ± 0.07 | -45.0 ± 2.5 | 2 |
| NaOAc<br>(37.5 mM)<br>+<br>Memantine<br>(30 μM) | 8 | -63.0 ± 3.4 | 39.0 ± 3.5 | 0.54 ± 0.10 | -47.5 ± 1.7 | 3 |
| NaOAc<br>(75 mM) | 7 | -59.9 ± 1.8 | 42.2 ± 1.6 | 0.40 ± 0.04 | -47.7 ± 1.1 | 3 |

Values are mean ± SEM; n = number size, $V_m$ resting membrane potential, $C_m$ capacitance, $HR_{input}$ hyperpolarizing input resistance, $V_t$ voltage threshold for action potential, NaOAc sodium acetate.

### Preparation of neuronal cultures

Whole brain neuronal cultures were made from 1-day old SD rats according to our previous publication.(20, 21) Briefly, rats were anesthetized with pentobarbital (50 mg/kg) and the brains were extracted and dissected. Dissected brain areas were combined, and brain cells were dissociated before being plated on poly-L-lysine precoated culture dishes. Two days following plating, cultures were treated for two days with cytosine arabanoside (ARC, 10 μM) to kill off rapidly dividing cells (astroglia and astrocytes) followed by fresh ARC free media for the remainder of cultures. Neuronal cultures contained > 90% neurons with the remainder primarily astroglia. Neuronal cell cultures were grown for 10-14 days before use in cell culture experiments.

### Real-time calcium imaging

Primary neuronal cultures 12-14 days post isolation were incubated for 30 min with Fluo-4AM (Thermofisher, 3 μM final concentration) according to the manufactures instructions in artificial cerebral spinal fluid (ACSF), containing (in mM): 125 NaCl, 2 KCl, 2 MgSO_4_, 1.25 NaH_2_PO_4_, 26 NaHCO_3_, 2 CaCl_2_, 10 d-glucose, and 0.4 ascorbic acid (osmolality: 295–302 mosmol L^-1^; pH 7.3-7.4) in a 5% CO_2_ humidified incubator at 37 °C. The staining solution was aspirated and fresh ACSF (0.5 mL) added to the cells. Baseline calcium fluorescence were taken followed by the addition of acetate and/or acetate and memantine in ACSF (0.5 mL additional volume added) to reach the desired concentrations (e.g., 0.5 mL ACSF initial, then add 0.5 mL acetate (75 mM) in ACSF. Final concentration acetate (37.5 mM)). Images were captured for each well in a 24 well plate using a cell imager equipped with the correct LED lights (ZOE, Bio-Rad). Fluorescence intensity was analyzed and quantified using ImageJ software(22) on 10-13 randomly selected cells for each treatment and corrected fluorescence intensity (CFIT) compared between treatment groups prior to adjusting backgrounds. CFIT was calculated based on individual background fluorescence within each well, prior to batch background normalization for image preparation. Duplicates of each treatment were compared, and all showed similar trends. Negative controls were run for ACSF alone on time course for alterations in baseline calcium and there was no significant change in baseline values compared to 25 minutes post ACSF only.

### Justification on doses/concentrations

To verify the effects of ethanol in the CeA, we selected a dose of ethanol (1.7 μmole or 100 nL of 17.15 M) which was a dose we found to yield a maximum response *in vivo* as previously described.(13) Previous studies of targeted brain delivery of ethanol to the CeA in rats (75-350 mg%, consisting of 8,000-10,000 nL infusions or a total dose of 0.761 μmole ethanol) (23) have been reported, and our maximum dose utilized is close to this dosing regimen. The following acetate doses were used for *in vivo* CeA microinjections: 0.1 and 0.2 μmole or 100 nL of 1 M and 2 M sodium acetate. Previous intraventricular infusions (0.8-2.8 μmole) of acetate have been reported (24) and we are within acceptable limits.

For our electrophysiology experiments, we utilized the current ethanol metabolism profile (roughly 90% of ethanol is metabolized to acetic acid/acetate) (25). That is to say, ethanol metabolism to acetic acid/acetate is a 1:1 conversion and apparent ethanol concentrations measured would in theory produce equivalent acetic acid/acetate concentrations (e.g., 50 mM ethanol should equal 50 mM acetate). We constructed a dose-dependent response curve to acetate and found an EC_50_ of 5.90 mM and an E_max_ of 37.5 mM. We utilized 37.5 mM for our electrophysiology studies because it yielded the most robust response.

### Computational modeling

The NR1/NR2A NMDAR (2A5T) crystal structure (26) was downloaded from the protein databank (PDB). Acetic acid, glutamic acid, glutaric acid, glycine and AP5 structures were downloaded from PubChem. All structures were imported into PyRx (27) and a docking box measuring (4.3×7.3×5.0 angstrom) was made around native glutamate. In a separate set of experiments, a docking box measuring (3.5×4.9×4.1 angstrom) was made around native glycine. Prior to computational docking, native glutamate or glycine was removed and each structure was docked for 8 iterations. Docking outputs were imported in PyMol(28) for display and ligand/receptor interaction purposes.

### Chemicals

All chemicals were obtained from Sigma-Aldrich (St Louis, MO, USA) with the exception of tetrodotoxin (Tocris Bioscience, UK) and tetraethylammonium sodium (Fluka BioChemika, Switzerland). Glacial acetic acid was obtained from Sigma-Aldrich (St. Louis, MO) and ultra-pure water (> 18 MΩ) obtained in-house by filtration and purification of house distilled water from an EasyPure II water filtration system (Barnstead).

### Statistical analysis

Data values were reported as mean ± SEM. Depending on the experiments, group means were compared using either paired, unpaired Student’s *t*-test, a one-way or a two-way ANOVA. Differences between means were considered significant at p < 0.05. Where differences were found, Bonferroni post hoc tests were used for multiple pair-wise comparisons. All statistical analyses were performed with a commercially available statistical package (GraphPad Prism, version 9.0).

## RESULTS

### Local metabolism of ethanol to acetic acid/acetate in the CeA activates NMDAR *in vivo*

Our lab previously reported that ethanol and acetate microinjected into the CeA of anesthetized rats resulted in a sympathoexcitatory effect and speculated that it was due to local metabolism of ethanol to acetate (13). Consistent with our previous findings, microinjection of ethanol into the CeA of anesthetized rats elicited a sympathoexcitatory and pressor response (Figure 1B,E,F). Because we speculated that this sympathoexcitatory response was due to local metabolism of ethanol to acetate, we co-applied ethanol with an alcohol dehydrogenase inhibitor (ADH), fomepizole (29) or an aldehyde dehydrogenase inhibitor (ALDH), cyanamide (30), into the CeA (Figure 1C,D). Blockade of ADH with fomepizole or ALDH with cyanamide significantly blunted the sympathoexcitatory response from ethanol (one-way ANOVA, p<0.0001, Figure 1), indicating that neither ethanol nor acetaldehyde was likely a causitive agent in the excitatory response. Summary data for change in SSNA and MAP in response to ethanol, ethanol + fomepizole or ethanol + cyanamide are displayed in Figure 1. Cyanamide (0.24 nmole) alone had no significant effect on SSNA (pre: 0.04 ± 0.009 vs post: 0.04 ± 0.008, p=0.3016, paired t-test), LSNA (pre: 0.015 ± 0.004 vs post: 0.020 ± 0.005, p=0.10, paired t-test) or MAP (pre: 113 ± 6.0 vs post: 104 ± 8.0, p =0.0734, paired t-test) (n=5 rats).

Next, we verified that acetate was the bioactive metabolite from ethanol metabolism. Microinjection of acetate into the CeA of anesthetized rats increased sympathetic nerve activity and mean arterial blood pressure (MAP) in a dose-dependent manner (Figure 2). For NMDAR antagonism, memantine has seen utilization in alcohol studies (31, 32) to prevent cognitive decline and as a possible therapeutic for excitotoxicity (33). Furthermore, it is an FDA approved NMDAR blocker for the treatment of Alzhiemer’s disease (34), and has been shown to prevent alcohol craving in humans (35). The use of memantine would be anticipated to give more powerful treatment options clinically compared to utilizing AP5 as previously described (13) or MK-801 (Olney’s lesions) (36). Blockade of NMDAR within the CeA using memantine or AP5 significantly attenuated the acetate induced sympathoexcitatory and pressor responses (one-way ANOVA, p<0.05, Figure 2A,B,C) suggesting that increases in SNA were due to acetate activation of NMDAR (Figure 2). Memantine (3.0 nmole) alone had no significant effect on baseline MAP (pre: 112 ± 7.63 vs post: 110.3 ± 8.67, p =0.58, paired t-test), SSNA (pre: 0.023 ± 0.008 vs post: 0.024 ± 0.009, p=0.37, paired t-test) and LSNA (pre: 0.018 ± 0.002 vs post: 0.019 ± 0.002, p=0.14, paired t-test) (n=3 rats). Likewise, our previous study utizling AP5 saw no significant difference to CeA microinjected AP5 on baseline parmeters (13).

**Figure 2.**
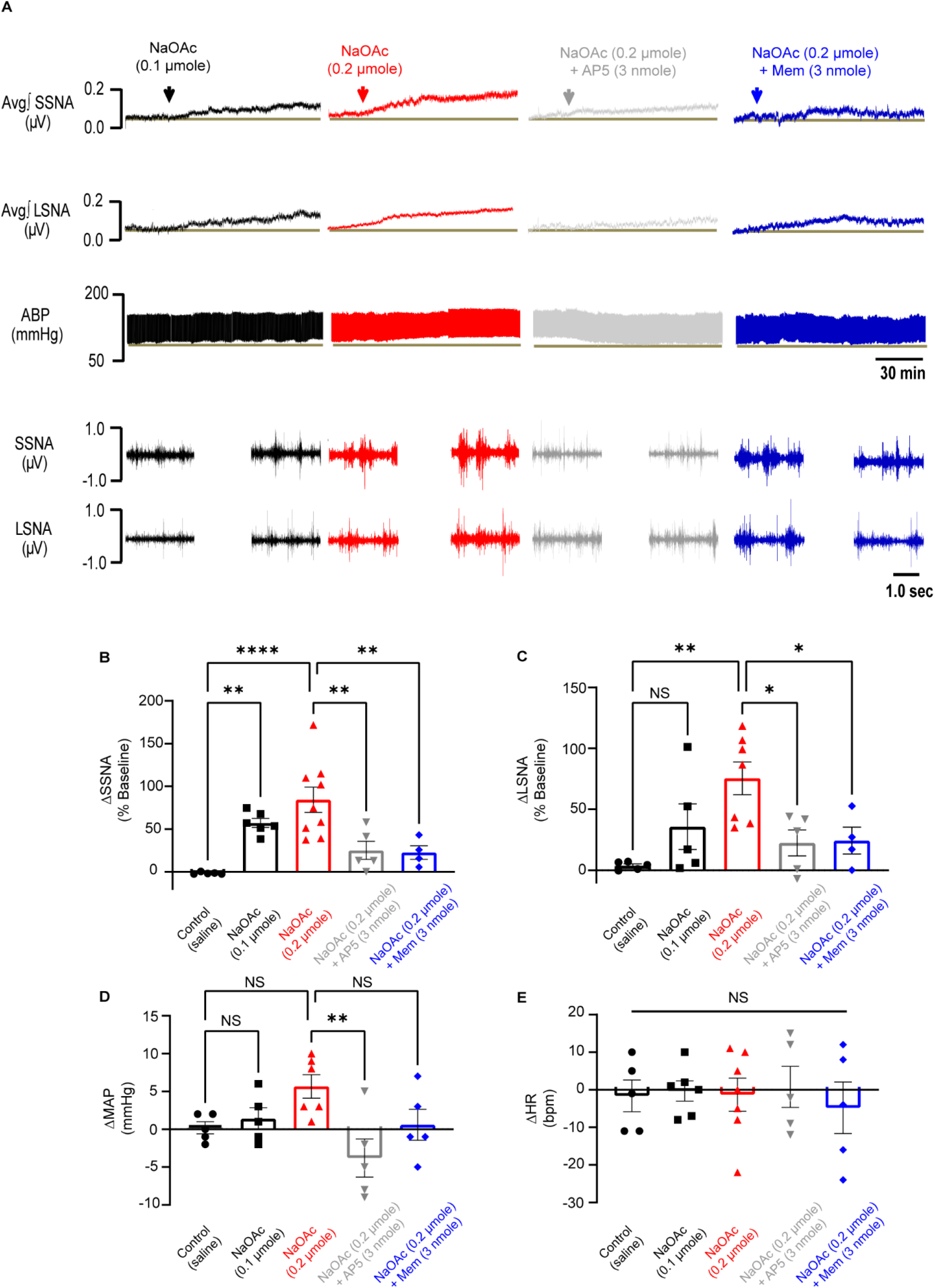
CeA microinjected acetate dose-dependently increases sympatheic nerve activity and is NMDAR dependent. **(A)** Representative raw traces displaying sympathetic nerve activity (SSNA and LSNA) and ABP from CeA microinjected acetate. **(B)** Summary SSNA data (**p=0.0093, **** p<0.0001). **(C)** Summary LSNA data (*p=0.045, **p=0.018). **(D)** Summary MAP data (**p=0.003). **(E)** Summary change in HR data (NS=not significant), (individual points=number of animals).

### Estimating final CeA tissue concentrations from microinjection

Estimation of brain microinjected compounds is a notoriously difficult task. Small volumes, effects of anesthesia, and blood flow drastically alter what local brain area tissues are exposed to in terms of final concentrations. The ethanol literature reports ~90% ethanol metabolism to acetic acid/acetate, and this conversion is 1:1 (25). Reported intoxicating blood alcohol concentrations in humans and rodents typically fall in the range of 17-100 mM (37–39), with measurements in the cerebellum of around 90 mM (40). However, brain slice patch clamp recordings sometimes utilize ethanol concentrations in excess of 200 mM (41–43). To estimate brain tissue concentrations to compounds tested, we microinjected 100 nL of ethanol or saline loaded with Chicago blue dye within the CeA, collected the stained tissue and recorded the weight. CeA estimated concentrations were determined by [x μmole compound/average tissue volume (mL)]. These concentrations represent the upper theoretical concentration limit and are likely an overestimate as diffusion is greater than the visualized stained tissue excised. Nevertheless, the upper theoretical concentration for 1.7 μmole ethanol is 283 mM, for 0.1 μmole acetate is 16.6 mM and for 0.2 μmole acetate is 33.3 mM (Figure 3).

**Figure 3.**
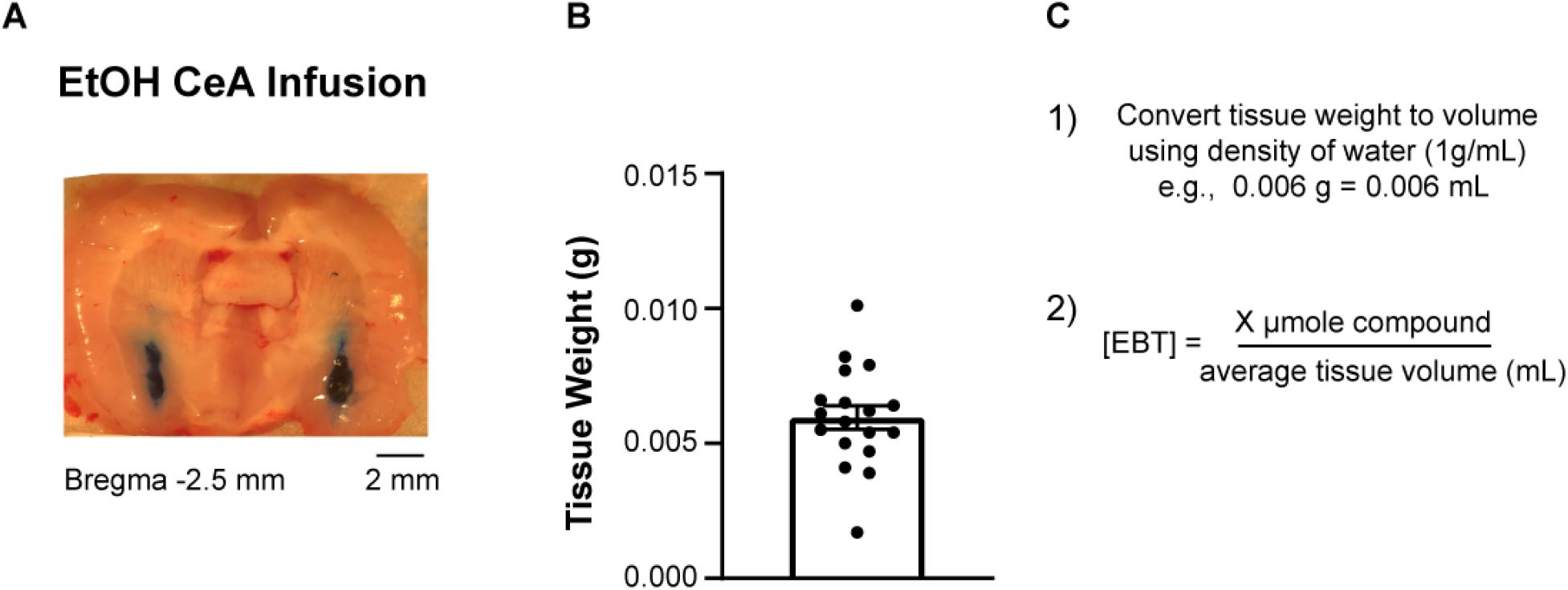
CeA brain tissue concentrations. **(A)** Representative image of CeA microinjection of 100 nL ethanol loaded with chicago blue dye. **(B)** Excised CeA tissue weights of microinjected ethanol or saline. **(C)** Method for estimating upper theoretical limit of brain tissue concentrations (n=18 CeA tissue samples from N=9 rats). Abbreviations: Estimated brain tissue = EBT.

### Bath application of acetate increases CeA-RVLM neuronal excitability through activatation of NMDAR and is not driven by excess extracellular sodium

Our previous *in vivo* microinjection study demonstrated ethanol and acetate induced sympathoexcitation by activation of NMDAR in the CeA through a projection to the rostral ventraolateral medulla (CeA-RVLM) (13). We therefore examined whether acetate was capable of increasing neuronal excitability of CeA-RVLM neurons in brain slice preparation. The RVLM as we and others have previously reported (13, 44, 45) is a key source of sympathoexcitatory drive that can contribute to elevated sympathetic outflow and heavily regulates cardiovascular function. What we found was that extracellular acetate consistently increased CeA-RVLM neuronal excitability in a dose-dependent manner (0.0 mM: 19 ± 1.2 Hz, n=7; 3.75 mM: 21.8 ± 1.6 Hz, n=5, 7.5mM: 36 ± 7.1 Hz, n=6; 37.5 mM: 41 ± 4.1, n=8; 75 mM: 44 ± 5.0, n=7), with an EC_50_ value of 5.90 mM (Figure 4) and an E_max_ of 37.5 mM. The EC_50_ value is close to peak acetate concentrations of ~4.60 mM measured by Wang and colleagues in ethanol and acetate infused rats (1) and is also similar to cytotoxic doses we observed in dopaminergic-like PC12 cells (14). Our E_max_ value of 37.5 mM was utilized to elicit maximum responses to both intrinsic excitability as well as NMDAR mediated inward currents. A one-way ANOVA with Bonferroni post-hoc test revealed differences (*F_(4,28)_* = 6.41, p=0.0009) in firing frequency observed between control vs 7.5 mM, 37.5 mM acetate and control vs 75 mM acetate.

**Figure 4.**
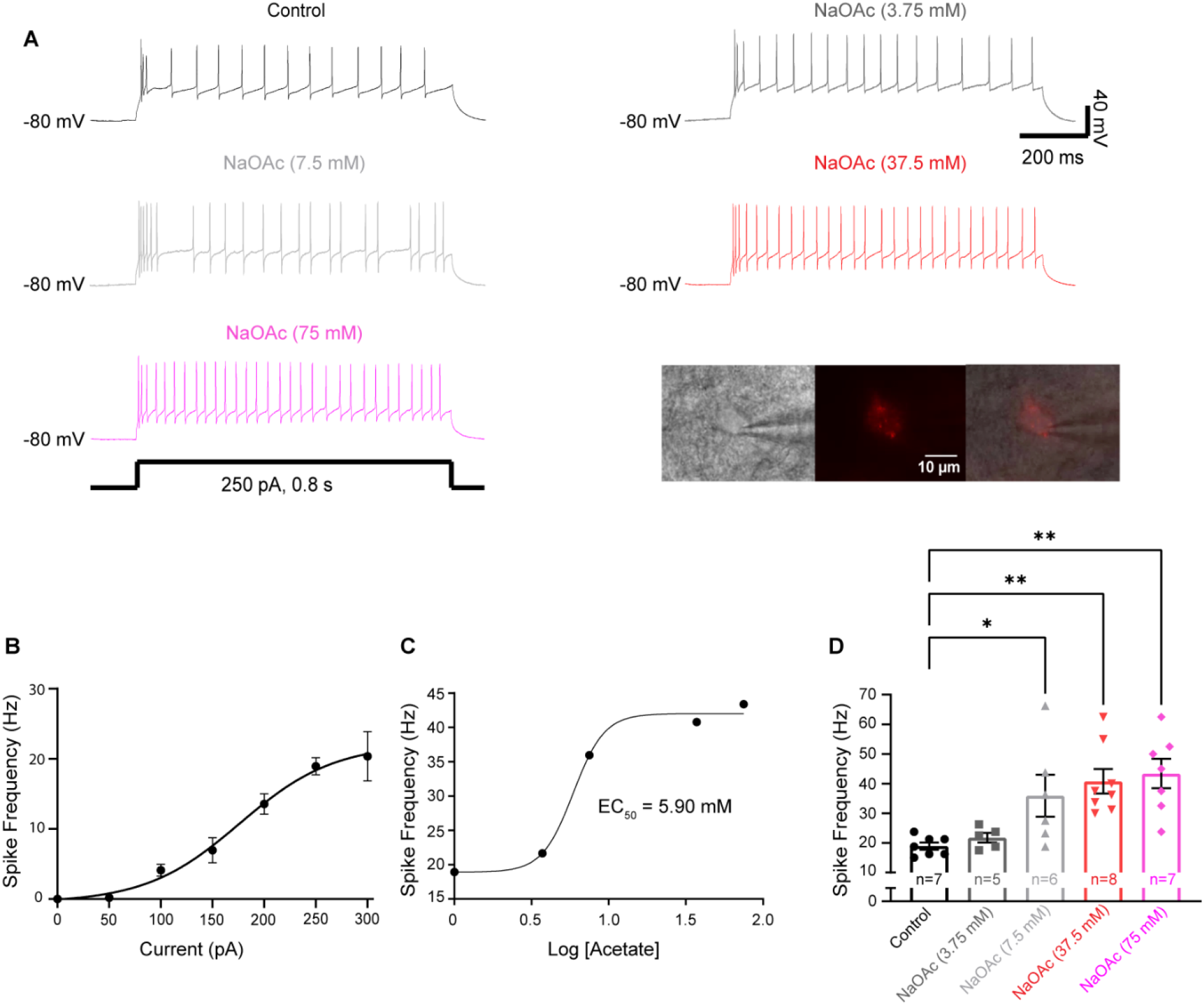
Acetate increases CeA-RVLM neuronal excitability in a dose-dependent manner. **(A)** Representative raw excitability traces for varying doses of acetate to +250 pA current injection and representative CeA-RVLM neuron; differential interference contrast (DIC, left), epifluorescence of the same neuron from retrograde RVLM fluorescent labeling (middle), merged DIC/epifluorescence indicating CeA neuron with projection to RVLM (right). Step-current injection protocol below traces. **(B)** Stimulus response curve for CeA-RVLM neurons (n=7 neurons, N=5 rats). Maximum firing occurred at +250 pA current injection. **(C)** Dose-dependent response curve for sodium acetate (NaOAc) on firing frequency at +250 pA current injection. The EC_50_ value for NaOAc was 5.90 mM. **(D)** Summary data for spike frequency at +250 pA current injection in control, and NaOAc (0, 3.75, 7.5, 37.5 and 75 mM) treated CeA-RVLM neurons (*p=0.0474, **p=0.0036, ***p=0.0016 vs control): (n=7 neurons, N=5 rats), (n=5 neurons, N=2 rats), (n=6 neurons, N=3 rats), (n=8 neurons, N=3 rats), (n=7 neurons, N=3 rats).

To exclude the possibility that increased excitability was driven by excess extracellular sodium concentrations and to verify that NMDAR were involved in the acetate induced increase in CeA-RVLM neuronal excitability, we used sodium gluconate (37.5 mM) as a sodium control and two NMDAR blockers, AP5 and memantine. Sodium gluconate (37.5 mM, N=2 rats, n=3 neurons) failed to increase neuronal excitability across the stimulus resopnse curve compared to control cells (two-way ANOVA, p>0.9999). Further analysis at 250 pA current injection also revealed no significant difference between control and sodium gluconate (19.00 ± 1.2 vs 15.00 ± 3.75 Hz, unpaired t-test, p=0.2225). Co-application of acetate and AP5 was only able to slightly reduce neuronal excitability (Figure 5A-C) while co-application of acetate and memantine (Figure 5A-C) was able to completely abolish the acetate induced increase in CeA-RVLM neuronal excitability. A two-way ANOVA with Bonferroni post-hoc correction of the current stimulus response (Figure 5B) showed significant differences among treatments (*F_(1,14)_* = 10.46, p= 0.0060). To understand the net effects of acetate on intrinsic excitability and the involvement of NMDAR over the range of the stimulus response, we analyzed the slope (gain) of each group’s curve (0-250 pA) using a linear regression. Linear regressions were well fit with R-squared values of (0.96, 0.95, 0.90, 0.95 and 0.94) for control, NaOAc, NaOAc & memantine and NaOAc & AP5, and sodium gluconate (37.5 mM) respectively. Acetate significantly increased the slope of the current injection response curve compared to control (0.157 ± 0.007 vs 0.066 ± 0.004 Hz/pA). Acetate and AP5 or acetate and memantine reduced the slopes to 0.1217 ± 0.005 Hz/pA and 0.0761 ± 0.008 Hz/pA respectively (Figure 5C). There was no statistical difference between control cells and sodium gluconate (0.066 ± 0.004 Hz/pA vs 0.06619 ± 0.008 Hz/pA, p>0.9999). A one-way ANOVA with Bonferroni post-hoc test revealed differences (*F_(3,24)_* = 47.81, p < 0.0001) between control vs acetate, acetate vs acetate + AP5, and acetate vs acetate + memantine. There was no statistical difference in passive neuronal properties highlighted in Table 1.

**Figure 5.**
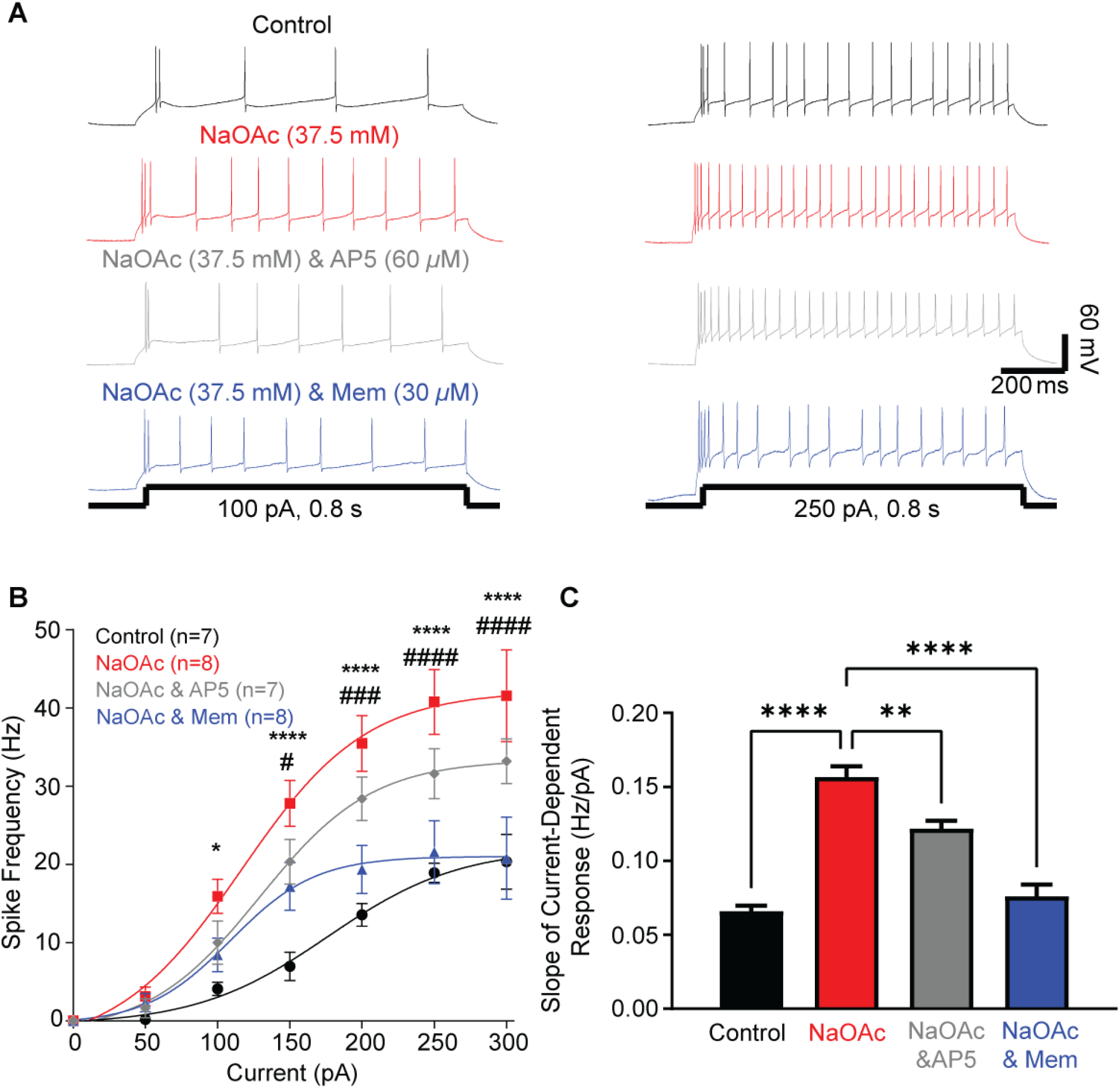
Effect of NMDAR antagonists on acetate induced excitability from current injection step protocol. **(A)** Representative raw traces for control (black), NaOAc (37.5 mM, red), NaOAc & AP5 (grey) and NaOAc & memantine (blue) at +100 pA (left) and +250 pA (right). Step-current injection protocols below traces. **(B)** Current stimulus response for control (black), NaOAc (red), NaOAc & AP5 (grey) and NaOAc & memantine (blue). (*p<0.05, ****p< 0.0001 NaOAc vs control, # p<0.05, *###* p<0.001, *###* p<0.0001 NaOAc vs NaOAc & memantine). **(C)** Slope of injected current response for control (black), NaOAc (red), NaOAc & AP5 (grey) and NaOAc & memantine (blue). (****p< 0.0001 vs control, ##p=0.0012, ####p< 0.0001 vs NaOAc) (n=7 neurons, N=5 rats), (n=8 neurons, N=3 rats), (n=6 neurons, N=2 rats) (n=6 neurons, N=3 rats).

### Computational modeling of glutamic acid, acetic acid, glutaric acid, glycine and AP5 with NR1/NR2A NMDAR

To help explain the discrepancy between AP5’s and memantine’s effectiveness in blunting the acetate-induced increase in excitability, we computationally modeled active glutamate site docking and binding affinity with 1) glutamic acid, 2) acetic acid 3) glutaric acid and 4) AP5 in the NR1/NR2A NMDAR(26) (Figure 6). Additionally, we modeled NR1/NR2A glycine site docking and binding affinity with 1) glycine and 2) acetic acid (Figure 6). Modeled structures were overlayed on the native glutamate ligand obtained from x-ray crystallography (2A5T) (26) or native glycine (2A5T). To demonstrate the effectiveness of computational modeling, we first modeled glutamic acid and compared it with natively bound glutamate. We found that computationally modeled glutamic acid orientated similarly to native glutamate (Figure 6) with similar ligand/receptor interactions and a binding affinity of −5.7 kcal/mol. Computationally modeled acetic acid orientated similarly to native glutamate carboxy terminals (Figure 6) with similar ligand/receptor interactions. The 1^st^ binding affinity near the glutamate carboxy location was −3.9 kcal/mol and the 2^nd^ binding affinity for the opposite carboxy terminal was −3.5 kcal/mol. Computationally modeled AP5 also orientated similarly to native glutamate (Figure 6) with similar ligand/receptor interactions and a binding affinity of −5.2 kcal/mol. To model potential binding of two acetic acid molecules simultaneously, glutaric acid was selected as it contains a five-carbon chain with free rotation about the chain’s axis and two carboxylic acid terminals (similar to acetic acid and similar to glutamic acid with no amine). Glutaric acid orientated similarly to native glutamate with similar ligand/receptor interactions and a binding affinity of −5.4 kcal/mol.

**Figure 6.**
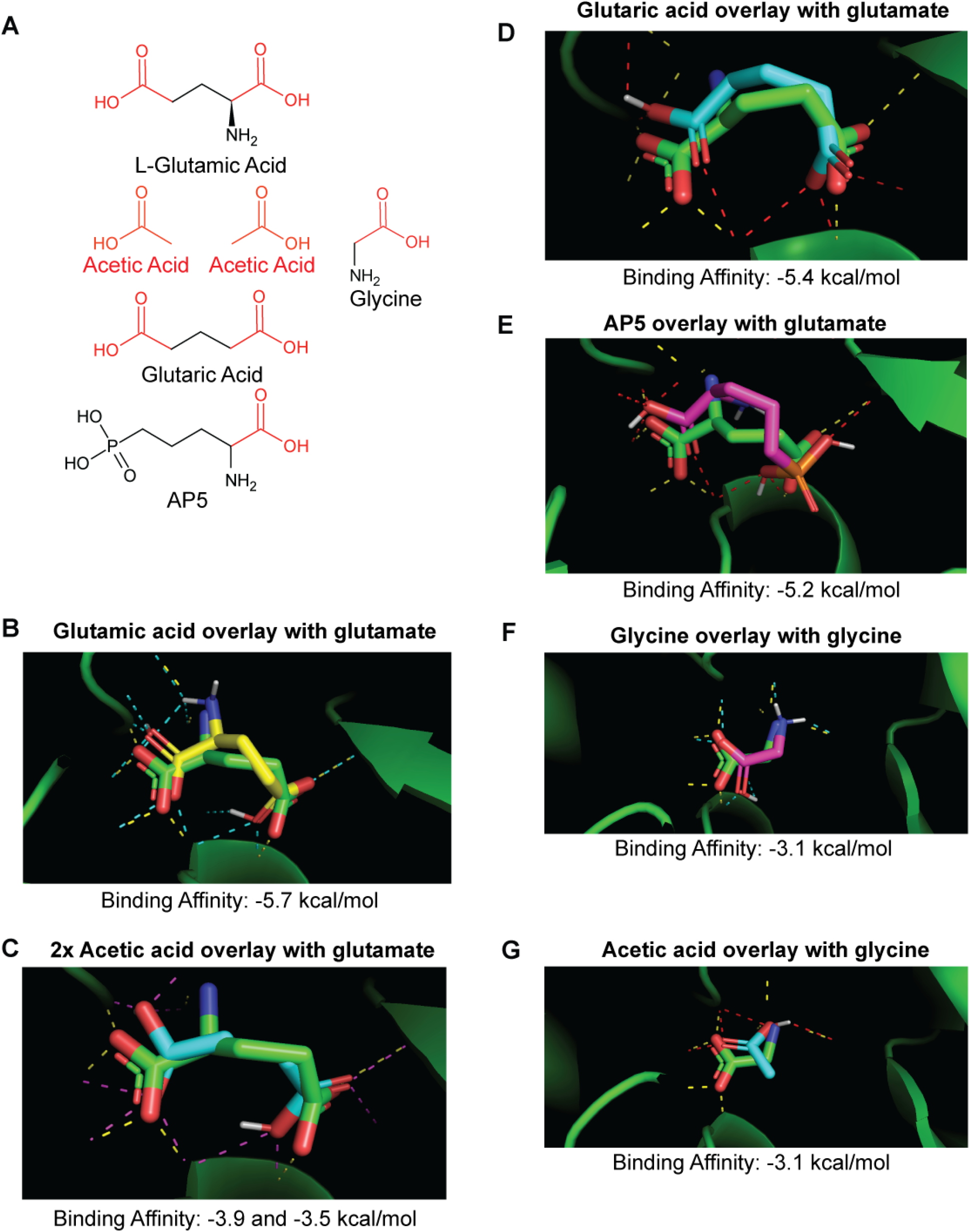
Computational modeling of NMDAR binding. **(A)** Structure of glutamic acid, acetic acid, glycine, glutaric acid and AP5. **(B)** Computational modeling of glutamate (yellow structure and interactions cyan) overlay in NR1/NR2A NMDAR with native glutamate (green structure and interactions yellow). **(C)** Computational modeling of two acetic acid molecules (cyan structure and interactions red) overlay in NR1/NR2A NMDAR with native glutamate (green structure and interactions yellow). **(D)** Computational modeling of glutaric acid (cyan structure and interactions red) overlay in NR1/NR2A NMDAR with native glutamate (green structure and interactions yellow). **(E)** Computational modeling of AP5 (magenta structure and interactions red) overlay in NR1/NR2A NMDAR with native glutamate (green structure and interactions yellow). **(F)** Computational modeling of glycine (magenta structure and interactions cyan) overlay in NR1/NR2A NMDAR glycine binding pocket with native glycine (green structure and interactions yellow). **(G)** Computational modeling of acetic acid (cyan structure and interactions red) overlay in NR1/NR2A NMDAR glycine binding pocket with native glycine (green structure and interactions yellow).

In the glycine binding pocket, computationally docked glycine had similar orientation to native glycine and similar ligand/receptor interactions (Figure 6) with a binding affinity of −3.1 kcal/mol. Computationally modeled acetic acid orientated slightly differently that native glycine but had similar ligand/receptor interactions and a calculated binding affinity of −3.1 kcal/mol.

### Bath application of acetate induces NMDAR mediated inward currents in CeA-RVLM neurons and is not driven by excess extracellular sodium

Next we examined whether external acetate application resulted in NMDAR mediated inward currents in CeA-RVLM neurons. We used memantine as our choice NMDAR antagonist because it showed greater efficacy in our intrinisic excitability protocols. What we found consistent with our excitability study was that bath application of acetate significantly increased NMDAR mediated inward currents, −22.5 ± 2.1 pA (Figure 7). Co-application of acetate and memantine was able to completely abolish NMDAR mediated inward currents, 7.3 ± 5.5 pA (Figure 7). Again, since we were applying acetate as a sodium salt, we wanted to verify that the increased NMDAR mediated inward currents were not due to increased sodium concentration. As such, we used sodium gluconate as a sodium ion control and found that 37.5 mM sodium gluconate was unable to trigger any NMDAR mediated inward currents, −1.1 ± 1.4 pA (Figure 7), suggesting acetate was the bioactive compound.

**Figure 7.**
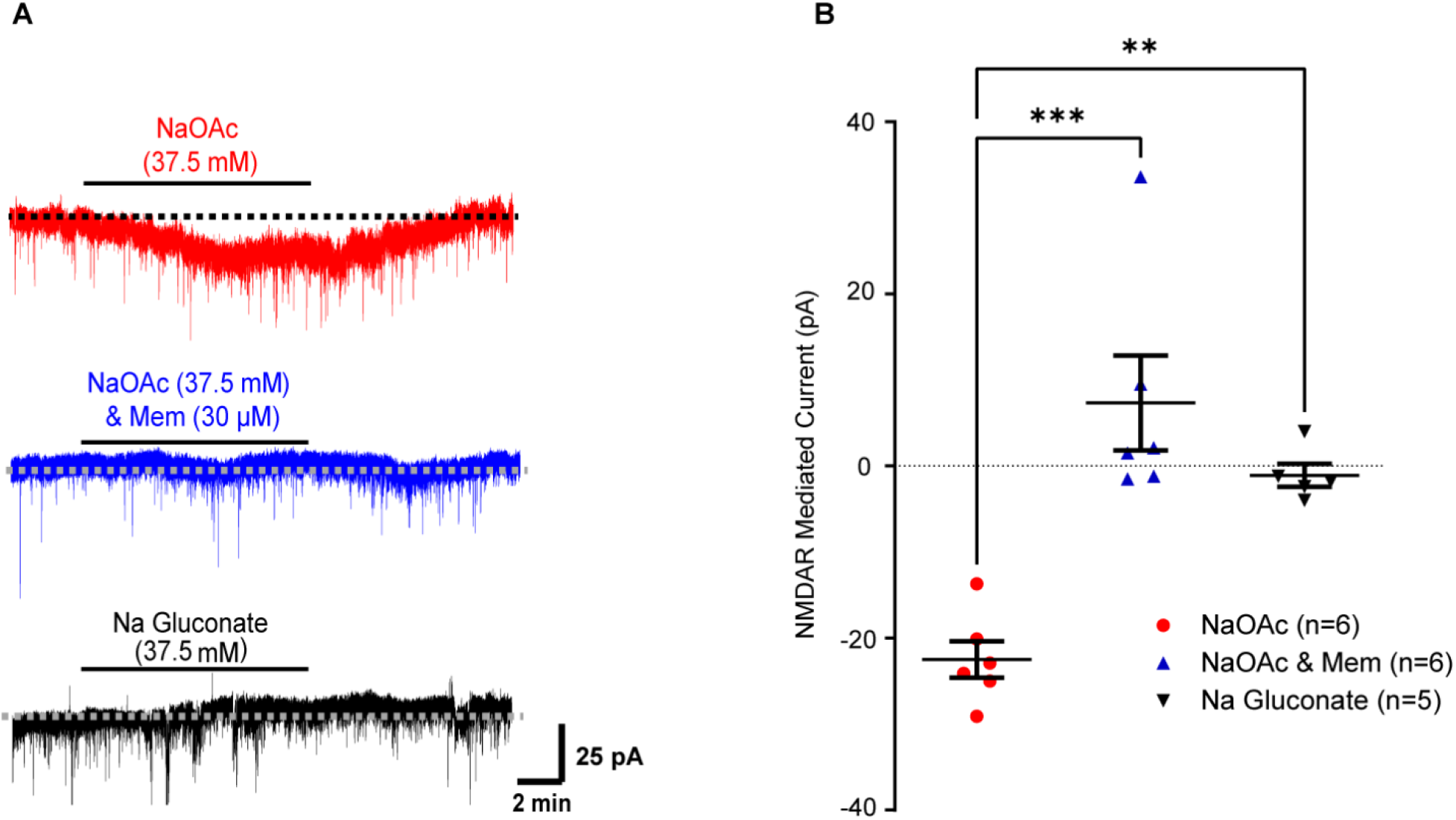
Acetate mediated NMDAR currents. **(A)** Representative raw traces for NMDAR mediated currents in CeA-RVLM neurons. **(B)** Summary data for NMDAR mediated currents in CeA-RVLM neurons. (***p=0.0001, **p=0.0038) (n=6 neurons, N=4 rats), (n=6 neurons, N=2 rats), (n=5 neurons, N=3 rats). Solid black lines above traces denote drug application time followed by washout

### Bath application of acetate increases depolarizing input resistance (R_input_) in CeA-RVLM neurons

To further understand the mechanistic effects of external acetate on CeA-RVLM neuronal excitability, neurons were subjected to a ramp protocol aimed to explore voltage threshold for firing an action potential (V_t_) and to determine depolarizing input resistance (Depolarizing R_input_) (Figure 8A-C). Input resistance (impedance) by definition is the measure of the opposition to current in an electrical network (ie, a greater depolarizing R_input_ suggests a greater resistance to positive current injection). Acetate increased the depolariing R_input_ compared to control (0.75 ± 0.04 GΩ vs 0.53 ± 0.05 GΩ) (Figure 8B), but had no statistically significant effect on alterations to V_t_ (Figure 6D). From an excitability standpoint, increased depolarizing R_input_ suggests an increase of positive ion influx in acetate treated neurons compared to control as they are depolarized. The effect of NMDAR blockers significantly reduced depolarizing R_input_, (*F_(3,25)_* = 4.00, P=0.0186) (Figure 8B).

**Figure 8.**
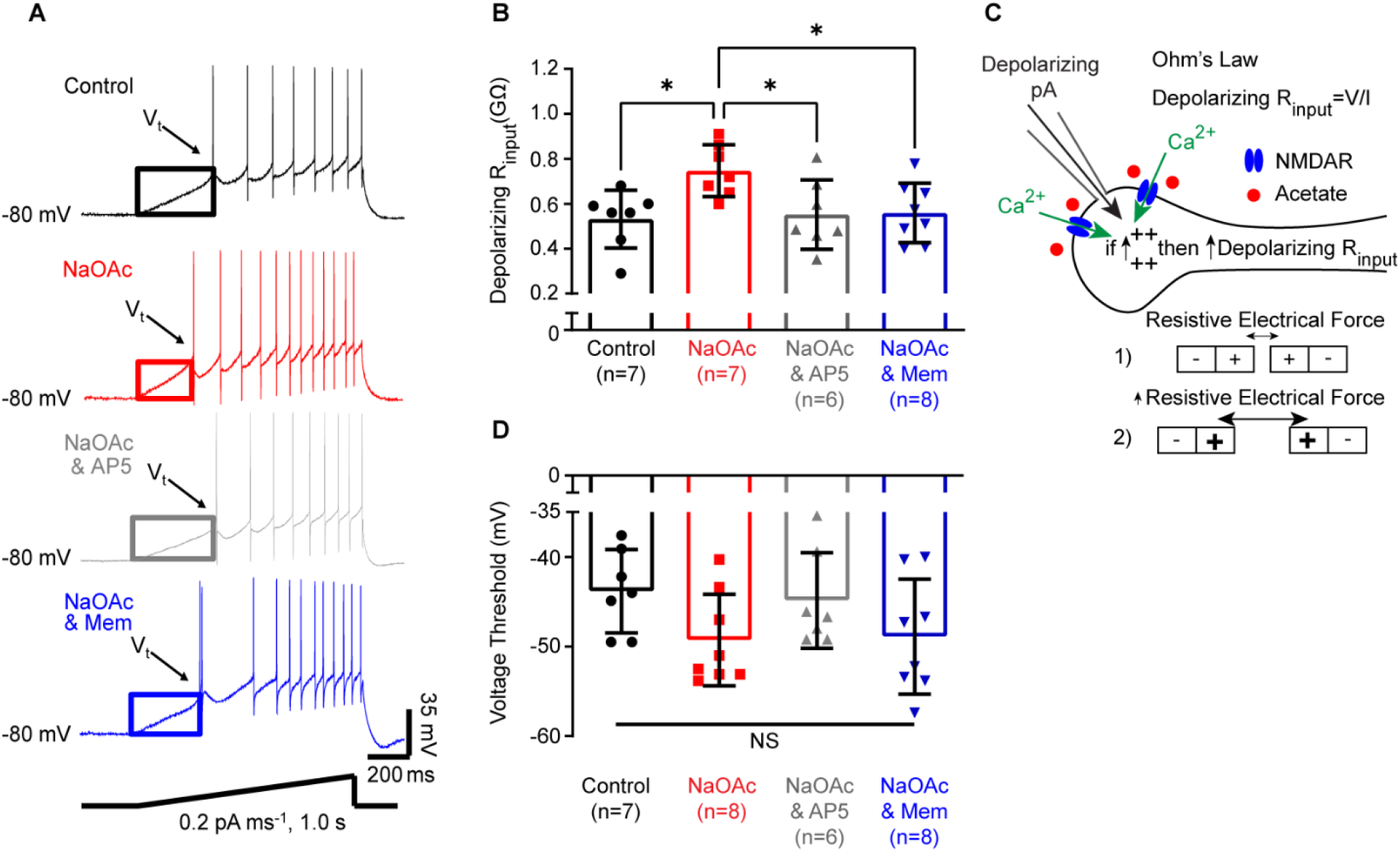
Effect of acetate on depolarizing input resistance (Depolarizing R_input_) and voltage threshold (V_t_) to firing an action potential. **(A)** Representative raw traces for ramp excitability protocol for control (black), NaOAc (red), NaOAc & AP5 (grey) and NaOAc & memantine (blue). Ramp-current injection protocol below traces. **(B)** Summary data for depolarizing R_input_. (* p < 0.05 vs control). **(C)** Graphical summary of depolarizing input resistance. Increased depolarizing current injection and activation of NMDAR results in an increase in positive charge buildup measured as a change in overall resistance via Ohm’s law. 1) Baseline electrical charge resistive force, 2) increased calcium enhanced electrical charge resistive force. **(D)** Summary data of voltage threshold to firing an action potential.

### Bath application of acetate to primary neuronal cultures increases cytosolic calcium and is NMDAR dependent

Our lab previously reported that bath application of acetate to dopaminergic-like PC12 cells increased cytosolic calcium (14) which was attenuated by co-application of acetate and memantine. We wanted to corroborate our previous findings in primary neuronal cultures. Bath application of acetate (37.5 mM, exact as those used in patch clamp recordings) increased cytosolic calcium fluorescence intensity in primary neurons in a time-dependent manner (baseline, 1 min and 25 min), 3,759 ± 430.5, 6,046 ± 500.7 and 6,775 ± 516.3 CFIT, respectively (*F_(2,36)_* = 10.58, p = 0.0002) (Figure 9 A-C, G). Co-application of acetate (37.5 mM) and memantine (30 μM) completely abolished any acetate induced increase in cytosolic calcium in neurons (*F_(2,27)_* = 0.21, p = 0.81) (Figure 9 D-F, H).

**Figure 9.**
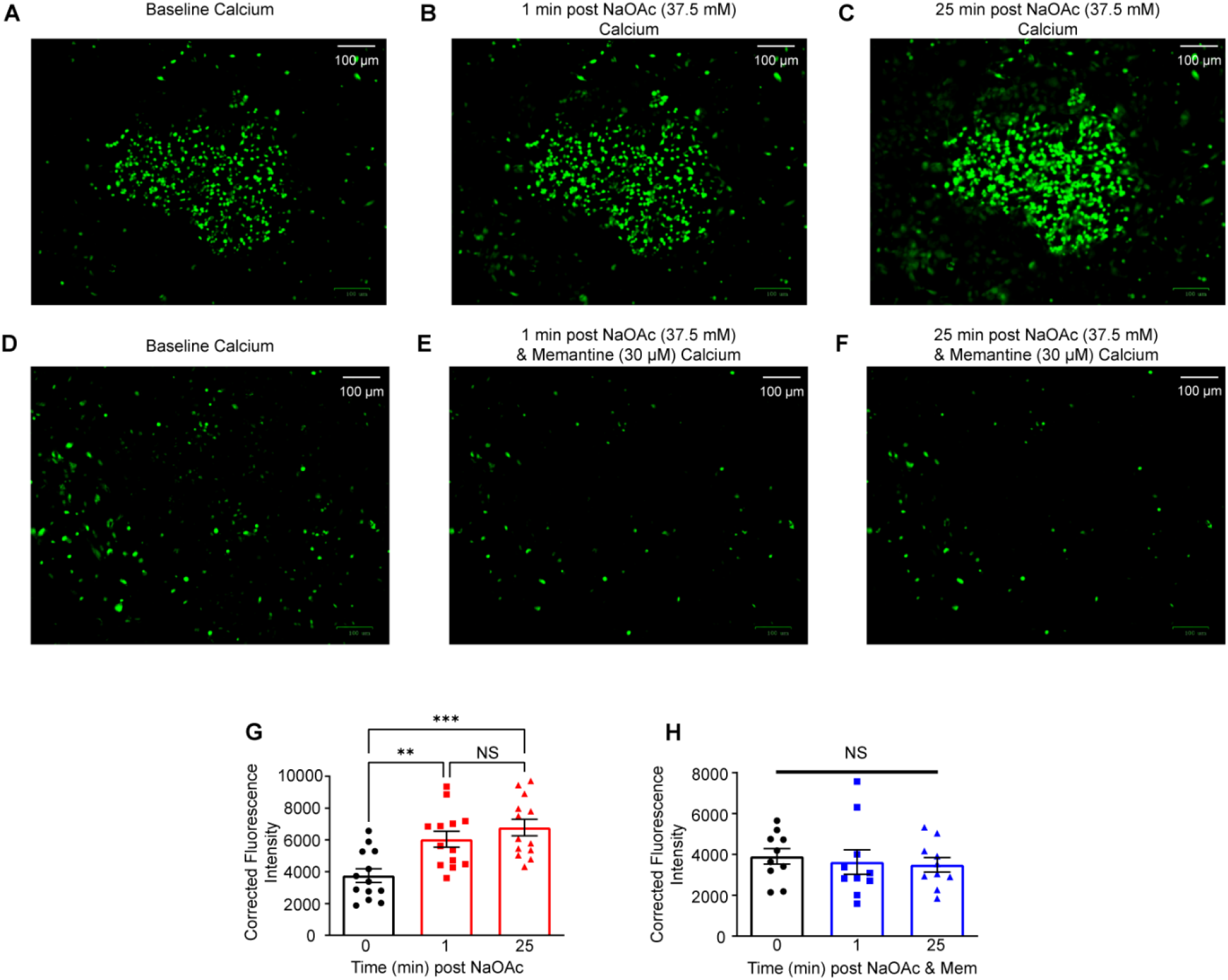
Acetate increases cytosolic calcium in neurons. **(A)** Baseline calcium fluorescence. **(B)** 1-minute post NaOAc. **(C)** 25 minutes post NaOAc. **(D)** Baseline calcium fluorescence. **(E)** 1-minute post NaOAc & memantine. **(F)** 25 minutes post NaOAc & memantine. **(G)** Neuronal calcium time-course summary data with NaOAc (**p<0.01, ***p<0.001 vs baseline). **(H)** Neuronal calcium time-course summary data with NaOAc & memantine (Mem). Note **A-C** and **D-F** images were acquired from different samples.

## DISCUSSION

In the present study, we evaluated how local brain metabolism of ethanol within the CeA elicits sympathoexcitatory responses *in vivo*. We find in a similar fashion to our previously reported work that microinjection of ethanol within the CeA increases SNA and MAP (13). Inhibiting local CeA ethanol metabolism with fomepizole (ADH inhibitor) or cyanamide (ALDH inhibitor) significantly blunts the ethanol induced sympathoexcitatory response. CeA microinjection of the ethanol metabolite, acetate, was found to increase SNA and MAP in a dose-dependent manner. These responses were attenuated by NMDAR antagonists, AP5 or memantine. In brain slice recordings containing CeA autonomic neurons with axon projecting to the RVLM (CeA-RVLM), bath application of acetate dose-dependently increased neuronal excitability with a calculated EC_50_ =5.90 mM and an E_max_ of 37.5 mM. Increased depolarizing input resistance was apparent in acetate treated CeA-RVLM neurons. Co-application of NMDAR antagonists attenuated these acetate induced responses, with memantine proving more effective. Similarly, NMDAR mediated inward currents in the presence of acetate were abolished by co-application with NMDAR antagonist. Finally, in primary neuronal cell cultures, bath application of acetate increased cytosolic calcium in a time-dependent manner, and this response was abolished by co-application of acetate and memantine. These findings suggest that ethanol-mediated inhibition of NMDAR *in vitro* is counteracted by acetate-mediated activation of the same receptor.

### Ethanol and neural control of cardiovascular function

Human studies have consistently demonstrated that acute ethanol consumption elicits sympathoexcitatory effects (46–48). While these effects are likely a combination of both peripheral and central mechanisms (49), centrally mediated mechanisms remain unclear. Furthermore, only a handful of studies have explored metabolic based mechanisms (13, 50). In our whole-animal SNA recordings from targeted brain delivery, we measure SSNA as well as LSNA. The SSNA closely corresponds with venous blood return and redistribution (51), while LSNA can be inferred as a close proxy for muscle SNA (52). While microinjection of acetate in the CeA increases both SSNA and LSNA, a key finding of increased LSNA may be linked to an effect noted in human studies. Acute ethanol was found to increase muscle SNA in humans (46, 47). Our results of CeA acetate on increased LSNA supports this previously mentioned finding in humans (46, 47), suggesting that increased muscle SNA in humans from ethanol may at least be partially mediated by centrally acting acetate. Future studies investigating the effects of microinjected acetate in other cardiovascular regulatory centers in the brain may delineate diverging, mechanistic and temporal output responses from acetate.

### Potential active site binding of acetate in NMDAR

Our electrophysiology studies demonstrated differences in the ability of two NMDAR antagonists to blunt the acetate-induced increase in neuronal excitability. We utilized the classic competitive NMDAR antagonist, AP5, and a non-competitive NMDAR pore-blocker, memantine. Our initial hypothesis regarding acetate activation of NMDAR was that the carboxy terminals of glutamate were partially agonized by two acetate molecules within the glutamate binding pocket (Figure 6). While memantine was able to completely abolish the response to acetate, AP5 was only partially effective. This suggests potential competition between AP5 and acetate for the active glutamate site in the NMDAR. We were able to corroborate our experimental findings with computational modeling and docking of glutamic acid, acetic acid and AP5 in the active site of NMDAR. Due to limitations in modeling we were unable to predict the binding affinity for two acetic acid molecules in the glutamate binding pocket. Our compromise was to select a compound with two terminal carboxylic acids and a non-ridgid carbon chain to allow for free rotation about its axis. Glutaric acid, an NMDAR agonist (53, 54), is a five carbon chain molecule with two terminal carboxylic acid functional groups (structurally similar to glutamate with no amine); it displayed higher simulated binding affinity compared to AP5 (−5.4 vs −5.2 kcal/mol), suggesting that two molecules of acetate may outcompete AP5. While we cannot definitively say that acetate actively binds at the glutamate binding pocket in NMDAR, our experimental findings and computational data support this possiblity.

We next modeled acetic acid at the glycine binding pocket in the NR1/NR2A NMDAR as acetic acid and glycine also share similar structural homology. Our computational data suggests that acetic acid has similar simulated binding affinity to glycine (−3.1 vs −3.1 kcal/mol). Agonism at the glycine site by acetate may also explain the effectiveness of memantine over AP5 at abolishing acetate-induced neuronal excitability, as glycine has been demonstrated to partially reverse AP5 blockade of NMDAR (55). Taken together, these simulated models suggest that future work investigating the effects of acetate at NMDAR and glycine receptors *in vivo* and *in vitro* are needed.

### Mechanisms for acetate induced increase in excitability within the CeA

While our present study suggests a major involvement of CeA NMDAR in the acetate induced effects on neuronal excitability and sympathoexcitatory responses, there are other potential contributing mechanisms. Acetate incorporation into glutamatergic (56, 57) as well as GABAergic (2, 40, 56) pools is likely a reflection of astrocytic metabolism. As such, astrocytic influence on glutamate cannot be ruled out. Previous work has demonstrated that acute acetic acid increased mEPSC frequency but not amplitudes, suggesting an increase in presynaptic glutamate (15). Whether this is from astrocytic generation or presynaptic synapses remains to be determined and has yet to be reported in CeA-RVLM neurons.

Another potential contribution to increased CeA excitability in the presence of acetate is AMPAR activation. We did see an ethanol induced AMPAR component in our previous microinjection study within the CeA (13), which is likely mediated via metabolism to acetate, however the NMDAR component appeared larger and thus we chose to focus our attention there. However, this AMPAR component may be a contributing factor seen in a lack of apparent effectiveness of AP5 and therefore we cannot rule out the impact of AMPARs in this response, especially as previous studies demonstrate an increase in mEPSC frequencies (likely AMPAR mediated events) in response to acute acetic acid in the accumbens shell (15). The fact that ethanol (44 mM) increased mEPSC frequencies with no apparent effect on intrinsic excitability supports the notion that this linkage to increased intrinsic excitability is not necessarily straightforward given the dissociation between the two (15). Nevertheless, these two previous studies demonstrate an inherent effect of acetic acid/acetate on AMPAR and this contribution deserves more attention in future studies.

### Perspectives

To the best of our knowledge, we are the first to report the effects of acetate/acetic acid on intrinsic exctiability of neurons, particularly autonomic CeA-RVLM neurons. As such, there is limited availability of literature with which we may compare our findings. Existing literature regarding acetate/acetic acid regulation of neuronal function seems supportive of the notion that both have robust effects in the CNS (2, 40, 58–61) which we also report here. Additionally, to the best of our knowledge there has only been one study describing the effect of ethanol/acetic acid interactions on neurophysiology (15). Future studies of these two pharmacological interactions should be pursued. Furthermore, given that NMDARs are heavily involved in synaptic plasticity (62) and alcohol use disorder (63), our finding that acetic acid/acetate activates NMDAR *in vivo* and *in vitro* may highlight a potential mechanism of alcohol use disorder pathogenesis. More studies are needed to determine the effects of centrally acting acetic acid/acetate on neuronal function and behavioral outputs, including cardiovascular function and whether there are any differences between sexes.

## Authors’ Contribution

A.D.C., M.J.H., K.M.D., J.E.B. and R.A.L. performed experiments. A.D.C., M.J.H., A.R.C., K.M.D., J.E.B., R.A.L., Z.S., L.Z. and Q.H.C. analyzed data. A.D.C., A.R.C., Q.H.C. and L.Z. prepared figures. A.D.C., A.R.C., Q.H.C. and L.Z. drafted the manuscript. All authors edited and revised manuscript. All authors approved the final version of the manuscript. A.D.C. and Q.H.C. conceptualized and designed the research.

## Acknowledgements

We would like to thank Dr. Timothy W. Chapp and Dr. Scott M. Chapp for their helpful suggestions and proof reading as well as Dr. Joe Erlichman and Dr. Ana Estevez for their early suggestions. The authors also thank Mrs. Mingjun Gu for excellent technical assistance.

